# Augmenting propulsion demands during split-belt walking increases locomotor adaptation in the asymmetric motor system

**DOI:** 10.1101/734749

**Authors:** Carly J. Sombric, Gelsy Torres-Oviedo

**Affiliations:** Department of Bioengineering, University of Pittsburgh, Pittsburgh, PA, United States

**Keywords:** Stroke, motor learning, hemiparesis

## Abstract

**Background:** Promising studies have shown that the mobility of individuals with hemiparesis due to brain lesions, such as stroke, can improve through motor adaptation protocols forcing patients to use their affected limb more. However, little is known about how to facilitate this process. Here we asked if increasing propulsion demands during split-belt walking (i.e., legs moving at different speeds) leads to more motor adaptation and more symmetric gait in survivors of a stroke, as we previously observed in subjects without neurological disorders.

**Methods:** We investigated the effect of propulsion forces on locomotor adaptation during and after split-belt walking in the asymmetric motor system post-stroke. To test this, 12 subjects in the chronic phase post-stroke experienced a split-belt protocol in a flat and incline session so as to contrast the effects of two different propulsion demands. Step length asymmetry and propulsion forces were used to compare the motor behavior between the two sessions because these are clinically relevant measures that are altered by split-belt walking.

**Results:** The incline session resulted in more symmetric step lengths during late split-belt walking and larger after-effects following split-belt walking. In both testing sessions, subjects who have had a stroke adapted to regain speed and slope-specific leg orientations similarly to young, intact adults. Importantly, leg orientations during baseline walking were predictive of those achieved during split-belt walking, which in turn predicted each individual’s post-adaptation behavior.

**Conclusion:** These results indicated that survivors of a stroke can adapt their movements to meet leg-specific kinetic demands. This promising finding suggests that augmenting propulsion demands during split-belt walking could favor symmetric walking in individuals who had a stroke, possibly making split-belt interventions a more effective gait rehabilitation strategy.

## Background

Brain lesions, such as stroke, may result in asymmetric gait (i.e., limp), limiting patients’ mobility and decreasing their quality of life (Jørgensen et al. 1995). Moreover, gait asymmetry can lead to comorbidities further affecting post-stroke gait such as musculoskeletal injuries (Jørgensen et al. 2000) and joint pain (Patterson et al. 2008). Promising studies show that split-belt walking, in which the legs move at different speeds, could correct gait asymmetries post-stroke (Reisman et al. 2007, 2009). However, it is not effective in all individuals (Reisman et al. 2013). It has been suggested that each subject’s baseline asymmetries are a factor limiting their ability to adjust their gait (Malone and Bastian 2014), raising the question of whether it would be possible to increase locomotor adaptation in this clinical population.

Our previous work indicates that locomotor adaptation in young, unimpaired subjects increases by augmenting propulsion demands during split-belt walking. More specifically, we found that baseline kinetic demands were predictive of step lengths at steady state and after-effects, such that greater propulsion demands led to more adaptation and larger after-effects in every individual (Sombric et al. 2019). It is unclear if the same could be observed post-stroke given their known propulsion deficits (Bowden et al. 2006; Balasubramanian et al. 2007). This might be possible since there is evidence that survivors of a stroke can augment their propulsion forces when required by the task (Kesar et al. 2011; Reisman et al. 2013; Awad et al. 2014; Hsiao et al. 2015, 2016b, 2016a). Thus, we tested whether locomotor adaptation in survivors of a stroke could be augmented by increasing propulsion demands with inclined split-belt walking.

We hypothesized that increasing propulsion demands would lead to more adaptation and after-effects following split-belt walking in individuals who have had a stroke. Thus, subjects in the chronic phase post-stroke experienced a split-belt adaptation protocol both in a flat and incline environment that had different propulsion demands (Lay et al. 2006, 2007). We expected more adaptation and greater after-effects following incline split-belt walking relative to flat split-belt walking. We also anticipated that those who have had a stroke would recover their baseline leg orientation to meet force demands at each inclination in the steady state split-belt condition. These anticipated findings would suggest that therapies increasing propulsion demands during walking would be a good strategy for improving post-stroke gait.

## Methods

We investigated the effect of augmenting propulsion demands during split-belt walking on gait adaptation under distinct slopes (i.e., flat and incline), which naturally modulate propulsion forces (Lay et al. 2006, 2007). To this end, we evaluated the adaptation and after-effects of 12 patients who have had a stroke (4 females, 61.1 +/− 10.6 years of age) in the chronic phase of recovery (>6 months post-stroke) during separate flat and incline testing sessions. Those who have had a stroke were eligible if they (1) had only unilateral and supratentorial lesions (i.e., without brainstem or cerebellar lesion) as confirmed by MRI, (2) were able to walk without assistance for 5 minutes at a self-selected pace, (3) were free of orthopedic injury or pain that would interfere with testing, (4) had no other neurological condition other than stroke, (5) had no severe cognitive impairments defined by a Mini-Mental State Exam score below 24, (6) could perform moderate intensity exercise, and (7) did not take medications that altered cognitive function. Written and informed consent was obtained from all participants prior to participation. The University of Pittsburgh Institutional Review Board approved the experimental protocol experienced by all participants.

### 2.1 General Paradigm

All subjects experienced a split-belt protocol while either walking flat or incline throughout two separate experimental sessions (Figure 1A). The flat session was always performed first. The protocol was tailored (i.e., slope, duration, and speed) so that each subject could complete both testing sessions at the same walking speed. The subject-specific walking speed on the treadmill was determined by subtracting 0.35 m/s from each subject’s overground walking speed during a Six-Minute Walking Test (Rikli and Jones 1998). We selected this procedure to ensure all individuals completed the entire split-belt walking protocol (Iturralde and Torres-Oviedo 2019). The walking speed, labeled as mid speed, for each participant is presented in Table 1. The speeds experienced during split-belt walking were selected based on subject’s mid walking speed. The slow speed was defined as 66.6% of the mid speed, and the fast speed as 133.3% of the mid speed. In this way, the average belt-speed during split-belt walking matched that of baseline and washout, and the belt-speed ratio during split-belt walking was 2:1. We selected an inclination of either 5° or 8.5° based on the level of the subject-specific motor impairments to ensure that all participants could complete the incline session.

**Table 1.**
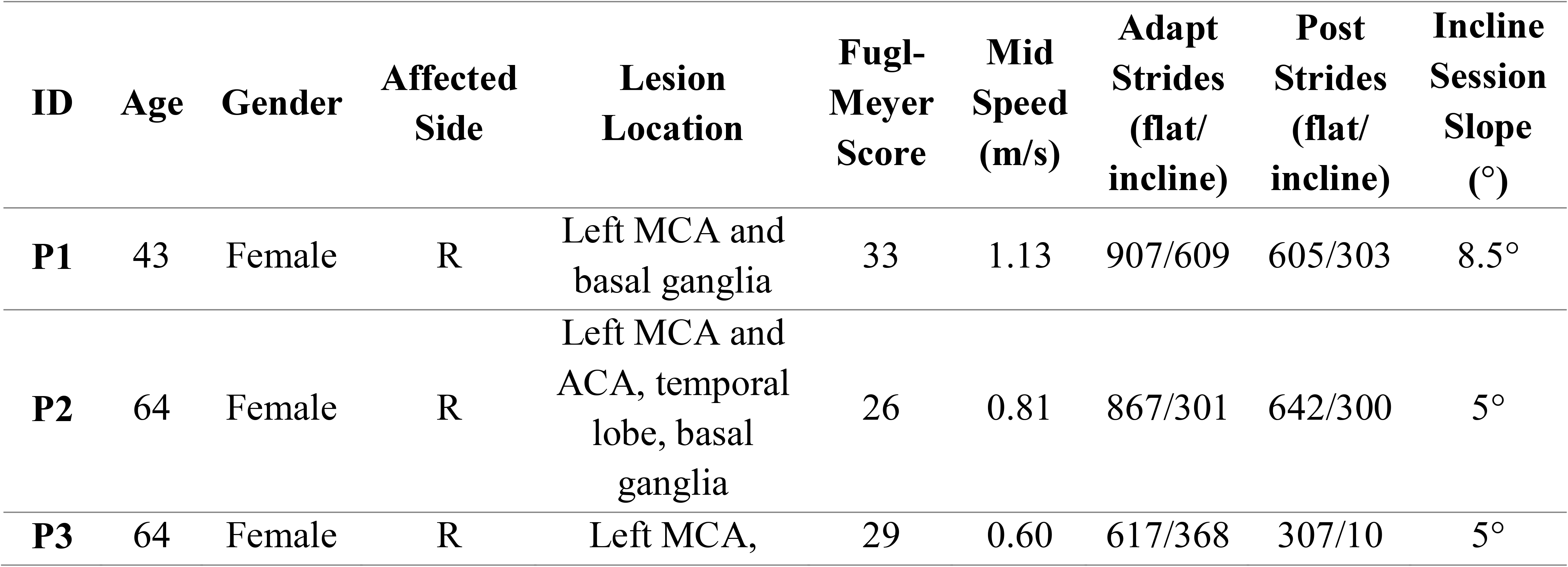
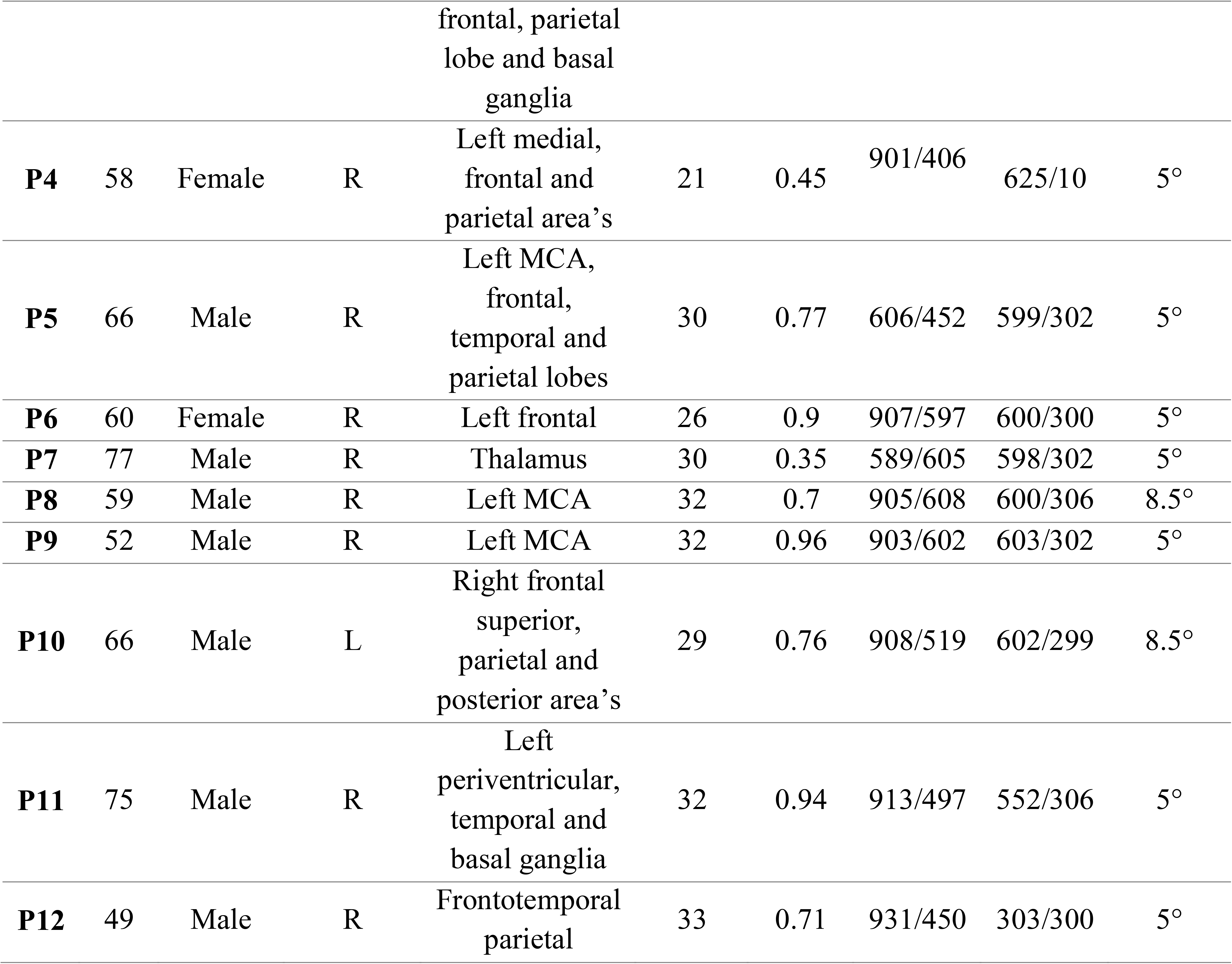
Clinical characteristics of stroke survivors.

| ID | Age | Gender | Affected Side | Lesion Location | Fugl-Meyer Score | Mid Speed (m/s) | Adapt Strides (flat/ incline) | Post Strides (flat/ incline) | Incline Session Slope (°) |
| --- | --- | --- | --- | --- | --- | --- | --- | --- | --- |
| P1 | 43 | Female | R | Left MCA and basal ganglia | 33 | 1.13 | 907/609 | 605/303 | 8.5° |
| P2 | 64 | Female | R | Left MCA and ACA, temporal lobe, basal ganglia | 26 | 0.81 | 867/301 | 642/300 | 5° |
| P3 | 64 | Female | R | Left MCA, frontal, parietal<br>lobe and basal<br>ganglia | 29 | 0.60 | 617/368 | 307/10 | 5° |
| **P4** | 58 | Female | R | Left medial,<br>frontal and<br>parietal area's | 21 | 0.45 | 901/406 | 625/10 | 5° |
| **P5** | 66 | Male | R | Left MCA,<br>frontal,<br>temporal and<br>parietal lobes | 30 | 0.77 | 606/452 | 599/302 | 5° |
| **P6** | 60 | Female | R | Left frontal | 26 | 0.9 | 907/597 | 600/300 | 5° |
| **P7** | 77 | Male | R | Thalamus | 30 | 0.35 | 589/605 | 598/302 | 5° |
| **P8** | 59 | Male | R | Left MCA | 32 | 0.7 | 905/608 | 600/306 | 8.5° |
| **P9** | 52 | Male | R | Left MCA | 32 | 0.96 | 903/602 | 603/302 | 5° |
| **P10** | 66 | Male | L | Right frontal<br>superior,<br>parietal and<br>posterior area's | 29 | 0.76 | 908/519 | 602/299 | 8.5° |
| **P11** | 75 | Male | R | Left<br>periventricular,<br>temporal and<br>basal ganglia | 32 | 0.94 | 913/497 | 552/306 | 5° |
| **P12** | 49 | Male | R | Frontotemporal<br>parietal | 33 | 0.71 | 931/450 | 303/300 | 5° |

**Figure 1:**
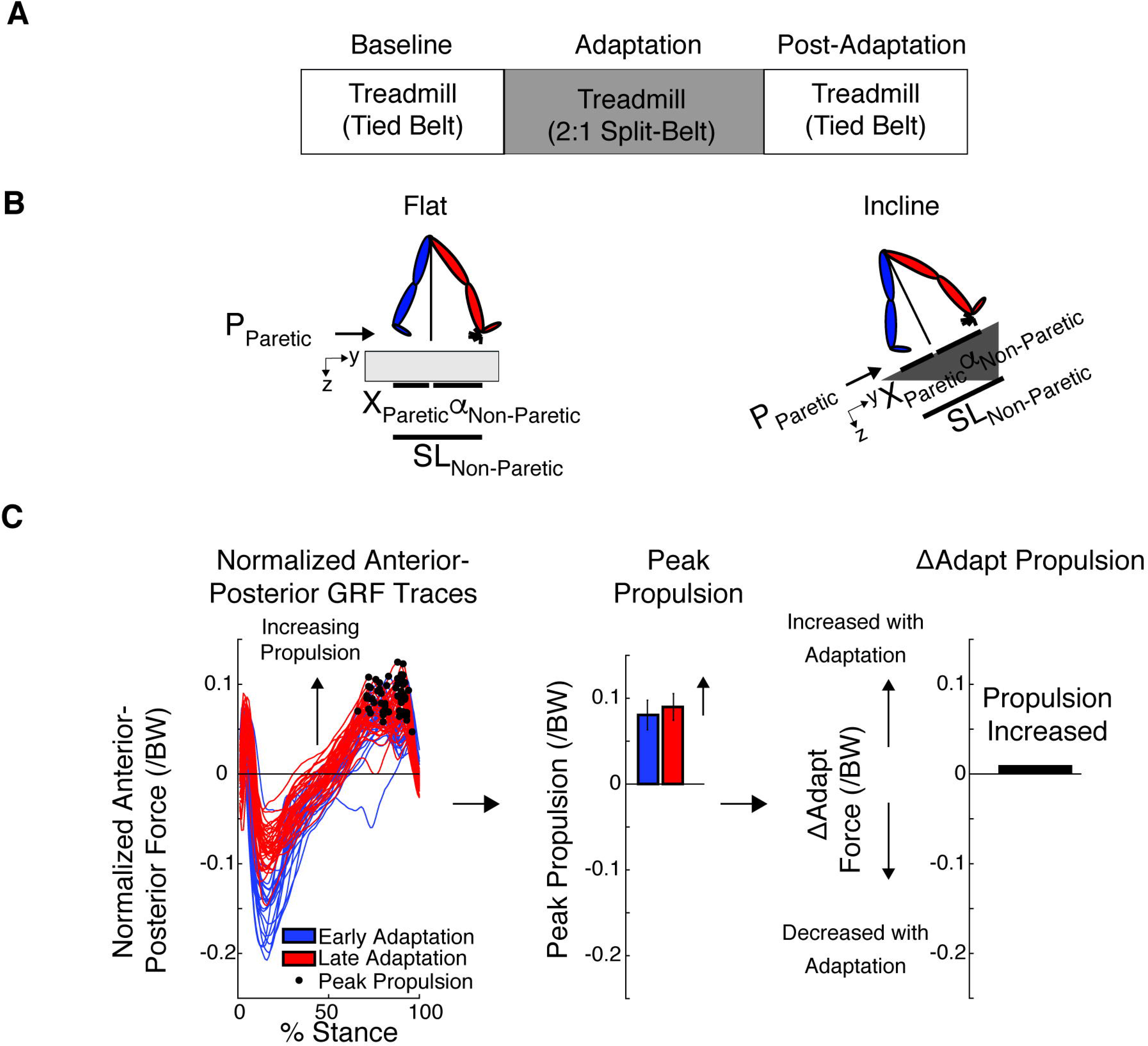
Experimental Paradigm and Kinetic and Kinematic Analysis. (A) Paradigm used for both the flat and incline sessions to assess locomotor adaptation during and after split-belt walking. Subjects walked flat for the entire flat session, and incline (either 5° or 8.5°) for the entire incline session. The walking speeds, duration of epochs, resting breaks and inclination were based on each subject’s ability. (B) The decomposition of step length into leading (α) and trailing (X) leg positions with respect to the body is illustrated for each sloped condition. This decomposition was done because it is known that inclination affects these aspects of step length differently (Leroux et al. 2002; Dewolf et al. 2017, 2018). Also note that when taking a step, the step length will depend on the position of the leading and trailing leg, which are generating a braking and propulsion force, respectively. (C) We used the peak propulsion force for each step to compute outcome measures of interest, such as the ΔAdapt measure. This measure was computed to quantify increments or reductions in magnitude within the adaptation epoch of each specific parameter. Note that increases in magnitude were defined as positive changes, whereas decreases in magnitude were defined as negative changes.

Experimental protocols for both sessions consisted of three epochs (i.e., Baseline, Adaptation, and Post-Adaptation). These epochs were used to assess subjects’ baseline walking characteristics and subjects’ ability to adjust and recalibrate their gait for each session-specific slope. Subjects first experienced a baseline epoch, lasting at least 50 strides, to characterize their baseline gait at the specific slope used throughout each session. Subjects walked with both belts moving at the same mid speed (Table 1). A baseline epoch with the belts moving at the slow walking speed (i.e., 66.6% of the mid speed) was also measured during the flat session. However, this epoch was removed in the incline session to ensure that all subjects could complete the entire protocol. Next, the Adaptation epoch was used to assess subjects’ ability to adjust their gait in response to a split-belt perturbation. During this epoch, the non-paretic leg walked twice as fast as the paretic leg. The paretic leg was confirmed with MRI. The duration of the Adaptation epoch for each subject are shown in Table 1. Finally, The Post-Adaptation epoch was used to assess the after-effects when the split-belt condition was removed. Both belts moved at the same mid speed as in the Baseline epoch. We counted the number of strides in real-time to regulate the duration for each epoch, where a stride was defined as the period between two consecutive heel-strikes (i.e., foot landings) of the same leg. All participants took resting breaks as requested. Also, all subjects wore a safety harness to prevent falls. In addition, there was a handrail in front of the treadmill for balance support, but individuals were encouraged to hold on to it only if needed.

### 2.2 Data Collection

Kinematic and kinetic data were used to characterize subjects’ ability to adapt their gait during Adaptation, and retain the learned motor pattern during Post-Adaptation. Kinematic data were collected with a passive motion analysis system at 100 Hz (Vicon Motion Systems, Oxford, UK). Subjects’ behavior was characterized with passive reflective markers placed symmetrically on the ankles (i.e., lateral malleolus) and the hips (i.e., greater trochanter) and asymmetrically on the shanks and thighs (to differentiate the legs). The origin of the kinematic data was rotated with the treadmill in the incline conditions such that the z-axis (‘vertical’ in the flat condition) was always orthogonal to the surface of the treadmill (Figure 1B). Gaps in raw kinematic data were filled with a quintic spline interpolation (Woltring; Vicon Nexus Software, Oxford Uk). Kinetic data were collected with an instrumented split-belt treadmill at 1,000 Hz (Bertec, Columbus, OH). Force plates were zeroed prior to each testing session so that each force plate’s weight did not affect the kinetic measurements. In addition, the reference frame was rotated at the session-specific inclination such that the anterior-posterior forces were aligned with the surface on which the subjects walked. A heel-strike was identified in real-time when the raw normal force under each foot reached a threshold of 30 N. This threshold was chosen to ensure accurate counting of strides at all slopes. During data processing we used a threshold of 10 N on median filtered data (with a 5 ms window) to detect the timing of heel strikes more precisely.

### 2.3 Data Analysis

#### 2.3.1 Kinematic Data Analysis

Kinematic behavior was characterized with step length asymmetry, which exhibits robust adaptation in split-belt paradigms (e.g., Reisman et al. 2005) and is of clinical interest. It is calculated as the difference in step length between the two legs on consecutive steps. Step length (SL) is defined as the distance in millimeters between the ankle markers at heel strike. Therefore, equal step lengths result in zero step length asymmetry. A positive step length asymmetry indicates that the non-paretic leg’s step length was longer than the paretic leg’s step length. Step length asymmetry was normalized by stride length, which is the sum of two consecutive step lengths, resulting in a unitless parameter that is robust to inter-subject differences in step size.

Each step length was also decomposed into anterior and posterior foot distances relative to the hip position (Figure 1B) as in previous work (Finley et al. 2015). This was done to quantify the leading and trailing legs’ positions relative to the body when taking a step because inclination is known to affect these measures (Leroux et al. 2002; Dewolf et al. 2017). The leading leg’s position (‘α’) was computed as the distance in millimeters between the leading leg’s ankle and the hip at heel strike; similarly, the trailing leg’s position (‘X’) was computed as the distance in millimeters between the trailing leg’s ankle and the hip at heel strike. The hip position, which is a proxy for the body’s position, was estimated as the mean instantaneous position across hip markers. By convention positive α values indicate that the foot landed in front of the hips, whereas negative X values indicate that the trailing leg was behind the hips. Note that the magnitudes of α and X summed to the leading leg’s step length. As indicated in Figure 1B, α and X were computed aligned to the treadmill’s surface in all sloped conditions.

#### 2.3.2 Kinetic Data Analysis

Kinetic data were used to characterize the adaptation of ground reaction forces (GRF). We focused our analysis on the propulsion component of the anterior-posterior GRF because they are associated with hemiparetic gait pathologies (Bowden et al. 2006; Balasubramanian et al. 2007). The anterior-posterior GRF (AP forces) were first low-pass filtered with a cutoff frequency of 20 Hz. Then, they were normalized by each subject’s body weight to account for inter-subject differences. Similar to our previous work, we computed peak propulsion forces (Sombric et al. 2019) as the maximum AP force (P_Paretic_ and P_Non-Paretic_) excluding the initial positive AP forces following heel strike. Note that we did not remove slope-specific biases due to gravity because we focused on analyzing changes in propulsion forces between epochs of interest.

#### 2.3.3 Kinetic and Kinematic Outcome Measures

Outcome measures were used to characterize kinematic and kinetic changes during the Adaptation and Post-Adaptation epochs relative to Baseline or within the Adaptation epoch. Outcome measures of interest were Baseline, Late Adaptation, After-Effects, ΔAdapt, and ΔPost. Baseline was defined as the average of the last 40 strides of the Baseline epoch for all parameters. This outcome measure characterized subjects’ baseline gait characteristics at each sloped environment and was used as a reference for Late Adaptation and After-Effects. Late Adaptation was defined as the difference between the average of the last 40 strides of the Adaptation epoch relative to Baseline for all parameters. This outcome measure indicated the steady state behavior reached at the end of the Adaptation epoch. After-Effects were defined as the difference between the average of the first 5 strides of Post-Adaptation and Baseline (e.g., Post-Adaptation - Baseline). Positive After-Effect values indicated increments in magnitude of a specific parameter during Post-Adaptation relative to Baseline, and vice versa for negative values. We also characterized the behavioral changes within Adaptation and Post-Adaptation with ΔAdapt and ΔPost, respectively. ΔAdapt was computed as the difference between Late Adaptation and Early Adaptation (i.e., average of the first 5 strides during the Adaptation epoch). ΔPost was computed as the difference between Baseline and early Post-Adaptation (e.g., Baseline - Post-Adaptation). Baseline was used instead of late Post-Adaptation because the duration of the Post-Adaptation epoch was not sufficiently long enough in all individuals to extinguish split-belt After-Effects. Thus, Baseline behavior was used a proxy for the late Post-Adaptation behavior. ΔAdapt and ΔPost were calculated such that an increase in the magnitude of a parameter resulted in positive values and vice versa. Figure 1C illustrates an example of the kinetic analysis for ΔAdapt.

### 2.4 Statistical Analysis

A significance level of α=0.05 was used for all statistical tests, which were performed either with Stata (StataCorp LP, College Station, TX) or with MATLAB (The MathWorks, Inc., Natick, Massachusetts, United States).

#### 2. 4. 1. Group analyses

Session averages were compared to determine the effect of slope on each of our outcome measures for kinetic and kinematic parameters. We tested the effect of slope on each one of our outcome measures (i.e., Baseline, ΔAdapt, ΔPost, Late Adaptation, After-Effects) with paired t-tests. In addition, we performed a one sample t-test to determine if ΔPost was significantly different from zero in the flat session.

During baseline walking, it was also of interest to identify differences between the paretic and non-paretic legs in addition to determining the effect of slope on outcome measures. Therefore, we performed ANOVAs with individual subjects as a random factor to account for the paired nature of the data set and slope and leg as fixed, repeated factors. These ANOVAs were performed on the peak propulsion values and the trailing leg’s position because our study was focused on the propulsion phase of the gait cycle, which is associated to these two parameters.

The changes of both step lengths during the Adaptation and Post-Adaptation epochs for the flat and incline sessions was also of interest. Therefore, we performed an ANOVA with individual subjects as a random factor to account for the paired nature of the data, and the fixed factors are slope, leg, and epoch. Slope and leg were considered repeated factors in the analysis. Epoch is not repeated and is treated as a between-subject factor given that these epochs are not directly associated (Sombric et al. 2019).

#### 2. 4. 2. Regression analyses

We tested the association between leg positions (‘α’ and ‘X’) during speed-specific Baseline and Late Adaptation to determine if Late Adaptation values could be predicted from Baseline values in survivors of a stroke, as observed in young unimpaired subjects (Sombric et al. 2019). We specifically tested the model |*y|* = *a_*_|z|*, where *y* is the predicted leg position during Late Adaptation and *z* is the leg position recorded during Baseline. We also tested the ipsilateral association between αs during Late Adaptation and Post-Adaptation and the contralateral association between Xs during these epochs in survivors of a stroke, since these relations were also observed in young, intact individuals (Sombric et al. 2019). Thus, we tested the model |*y|* = *a_*_|z|*, where *y* is each leg’s position during Early Post-Adaptation and *z* is either the ipsilateral ‘α’ position recorded during Late Adaptation or the contralateral ‘X’ position recorded during Late Adaptation. An absolute value of *z* was utilized so that the data would not bias the results of the regression to be linear by having a cluster of positive (α) and negative (X) data points.

## Results

### Adaptation and recalibration of step length asymmetry are augmented when walking incline

Step length asymmetry adaptation and recalibration were augmented by incline walking. Figure 2A illustrates the evolution of step length asymmetry throughout the flat and incline sessions. Figure 2B indicates that there was a wide range of individual Baseline step length asymmetries (colored lines) and slope did not change the group average biases (p=0.30). During Adaptation, participants exhibited similar changes in step length asymmetry from early to late Adaptation (Figure 2D, p=0.75), but they were more symmetric in the incline than the flat session in Late Adaptation (Figure 2C, p=0.004). Furthermore, the incline session had larger magnitudes of After-Effects during early Post-Adaptation relative to the flat session (Figure 2E, p=0.008). Thus, incline walking augmented step length symmetry during Late Adaptation and the magnitude of After-Effects.

**Figure 2:**
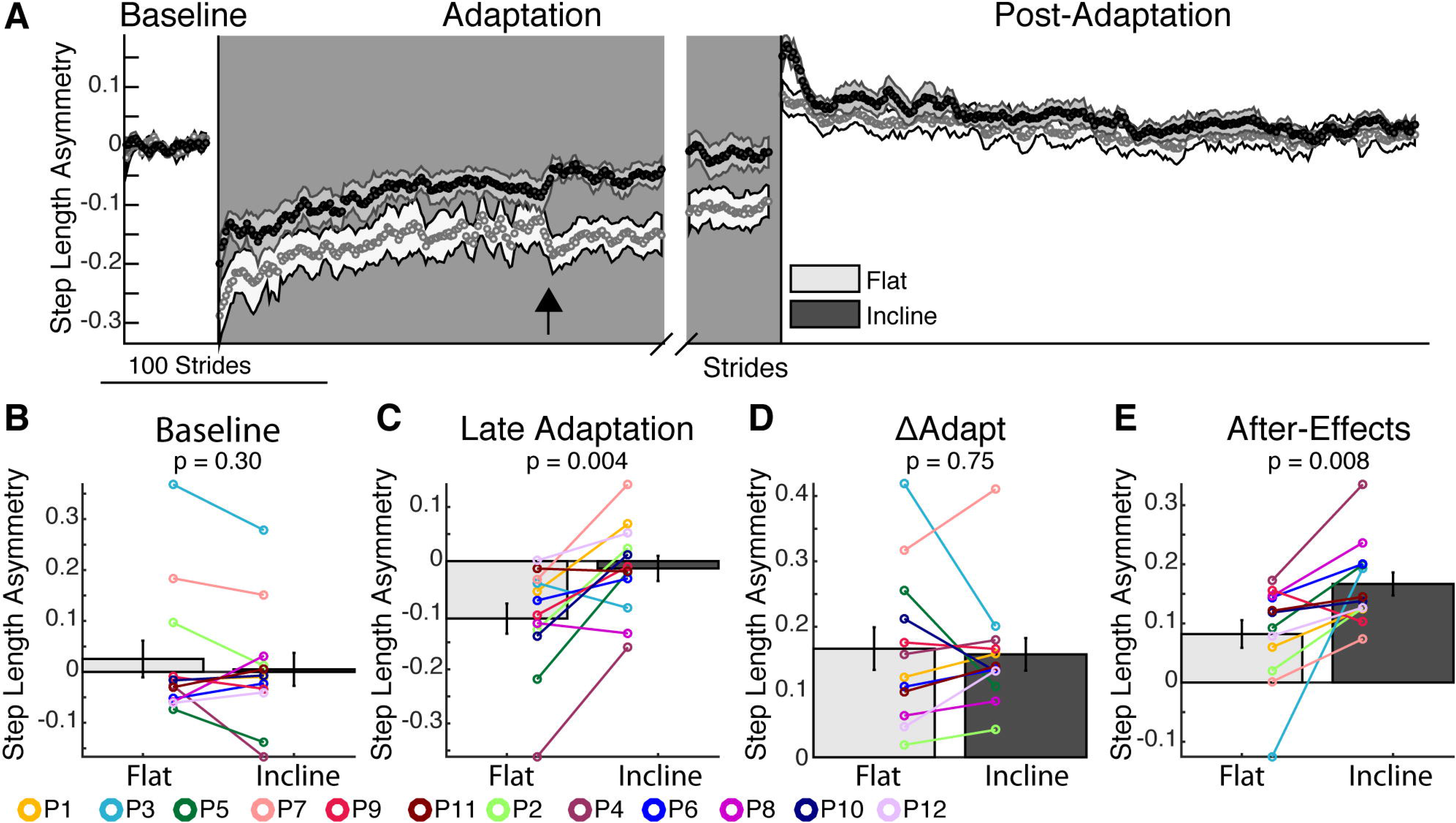
Step Length Asymmetry Adaptation and Recalibration. **(A)** Stride-by-stride time course of step length asymmetry during Baseline, Adaptation, and Post-Adaptation for each session are shown. Note that each subject’s baseline bias has been removed. Each data point represents the average of 5 consecutive strides and shaded regions indicate the standard error for each session. For display purposes only, we include in the time courses stride values that were computed with a minimum of 10 subjects. The black arrow indicates a discontinuity in the data caused by many subjects taking a resting break at the same time. **(B-E)** The height of the bars indicates group average step length asymmetry ± standard errors. Individual subjects are represented with colored dots connected with lines. **(B)** Baseline: Baseline step length asymmetry is not influenced by slope. **(C)** Late Adaptation: Note that each session plateaued at different step length asymmetry values during the Adaptation epoch such that subjects reached more symmetric step lengths in the incline session than the flat session **(D)** ΔAdapt: Participants changed their gait by similar amounts during the Adaptation epoch in both sessions. **(E)** After-effects: Subjects had larger After-Effects during early Post-Adaptation in the incline session than the flat session, which is consistent with the Late Adaptation differences across sessions.

### Both step lengths contribute to step length asymmetry adaptation and after-effects during incline walking in the asymmetric motor system

Survivors of a stroke adjusted both step lengths during split-belt walking. The survivors of a stroke modulate both their slow (paretic) and fast (non-paretic) step lengths during Adaptation and have After-Effects during Post-Adaptation (Figure 3A). The change of each step length during the Adaptation and Post-Adaptation epochs are quantified in Figure 3B. There was a significant effect of epoch (p_epoch_=0.001) and interaction between leg and epoch (p_leg#epoch_<0.001) indicating that the step length with the paretic leg is reduced during Adaptation, but increased during Post-Adaptation and vice versa for the non-paretic leg. Overall, slope did not alter step length changes (p_slope_=0.16, p_slope#leg_=0.18, p_slope#epoch_=0.17), except for the paretic leg’s de-adaptation as quantified by ΔPost (p_leg#epoch#slope_=0.016). More specifically, the paretic step lengths did not exhibit de-adaptation in the flat session (i.e., ΔPost is not different from zero, p=0.38), whereas step lengths for both legs had significant de-adaptation in the incline session (i.e., non-zero ΔPost, p<0.001). Survivors of a stroke use both their paretic and non-paretic leg to counteract the split-belt perturbation and both legs are recalibrated following incline adaptation.

**Figure 3:**
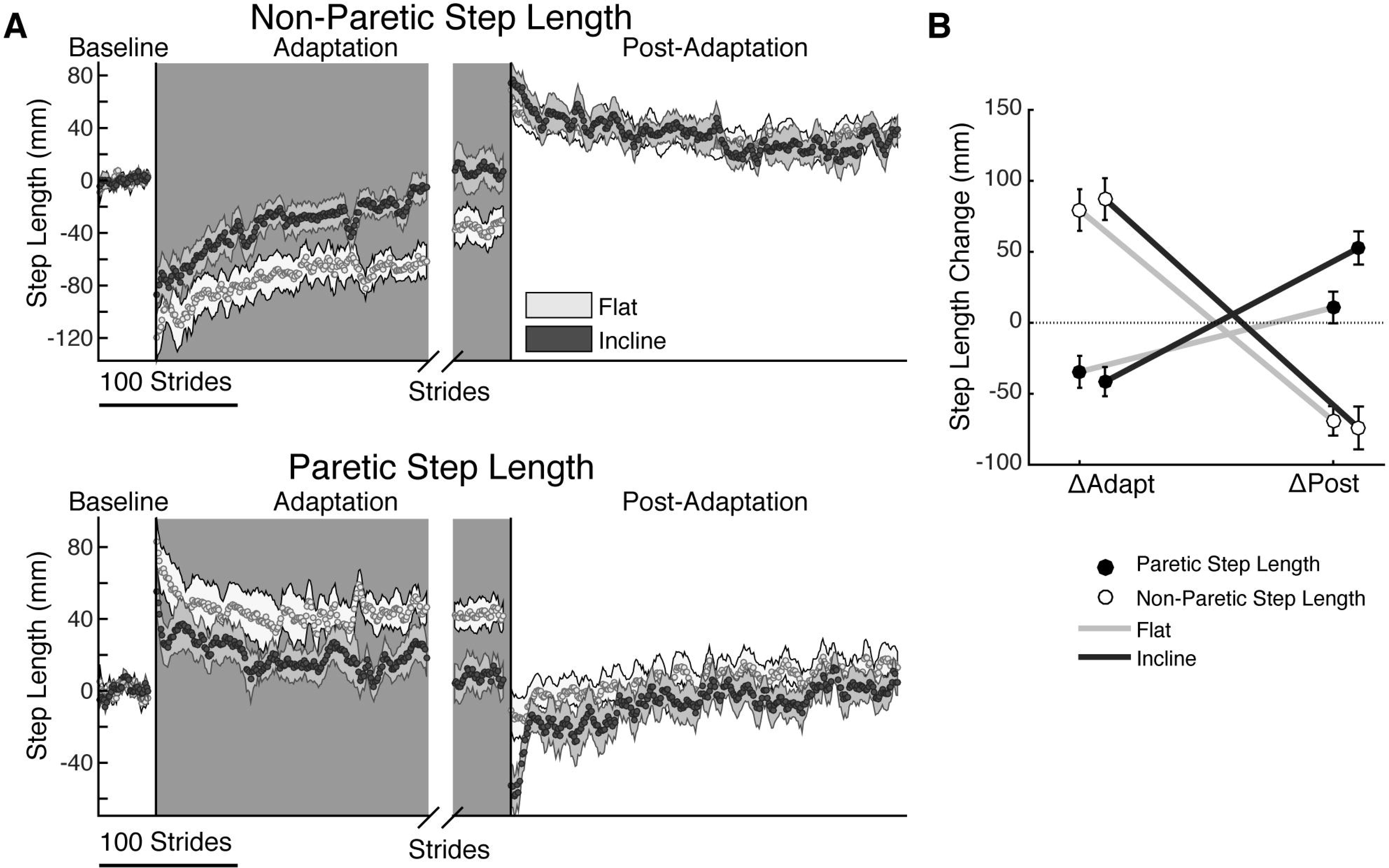
Step length Adaptation and After-Effects. **(A)** Stride-by-stride time courses of step lengths when either the non-paretic leg (top panel, fast leg during Adaptation) or the paretic leg is leading (bottom panel, slow leg during Adaptation) are shown during Baseline, Adaptation, and Post-Adaptation. Each data point represents the average of 5 consecutive strides and shaded regions indicate the standard error for each group. For display purposes only, we include stride values during Post-Adaptation that were computed with a minimum of 10 subjects. **(B)** The effect of slope on each leg’s change during Adaptation (ΔAdapt) and Post-Adaptation (ΔPost) is illustrated. Note that both the paretic and non-paretic leg adapted similarly. While the non-paretic leg has recalibrated (ΔPost≠0) following both the flat and incline session, the paretic leg is only recalibrated following incline Adaptation.

### Slope and speed-specific walking demands determine the distinct step length asymmetries across inclination conditions

Speed and slope-specific leg orientations mediated the distinct step length asymmetries selected during Late Adaptation and early Post-Adaptation. Figure 4A illustrates a top-down view of the baseline leg orientations that contribute to each step length relative to the hips. While we found significantly different leg orientations across individuals (colored lines, p_Individual_=0.002), we observed that the trailing leg position, X, was larger in the incline than flat condition for both legs (p_Slope_=0.042, p_Leg_=0.22, p_Slope#Leg_=0.76). The schematic in figure 4B illustrates the relation between baseline speed-specific leg orientations and late Adaptation (Sombric et al. 2019). Figure 4C indicates that the participants’ leg orientations during slow Baseline walking predict well those achieved during late Adaptation (solid cyan line; |y|=a*|x|; 95% confidence interval of a=[0.92, 1.13], R^2^=0.76, p<0.001). We also show as a reference, the relation between (recorded) Baseline and (predicted) Late Adaptation leg orientation values for both legs and both inclinations in young unimpaired individuals (Sombric et al. 2019) (magenta dashed line; (|y|=a*|x|; 95% confidence interval of a=[0.91, 0.96], R^2^=0.89, p<0.001). Note the similarity between the intact and lesioned behavior (cyan vs. magenta lines).

**Figure 4:**
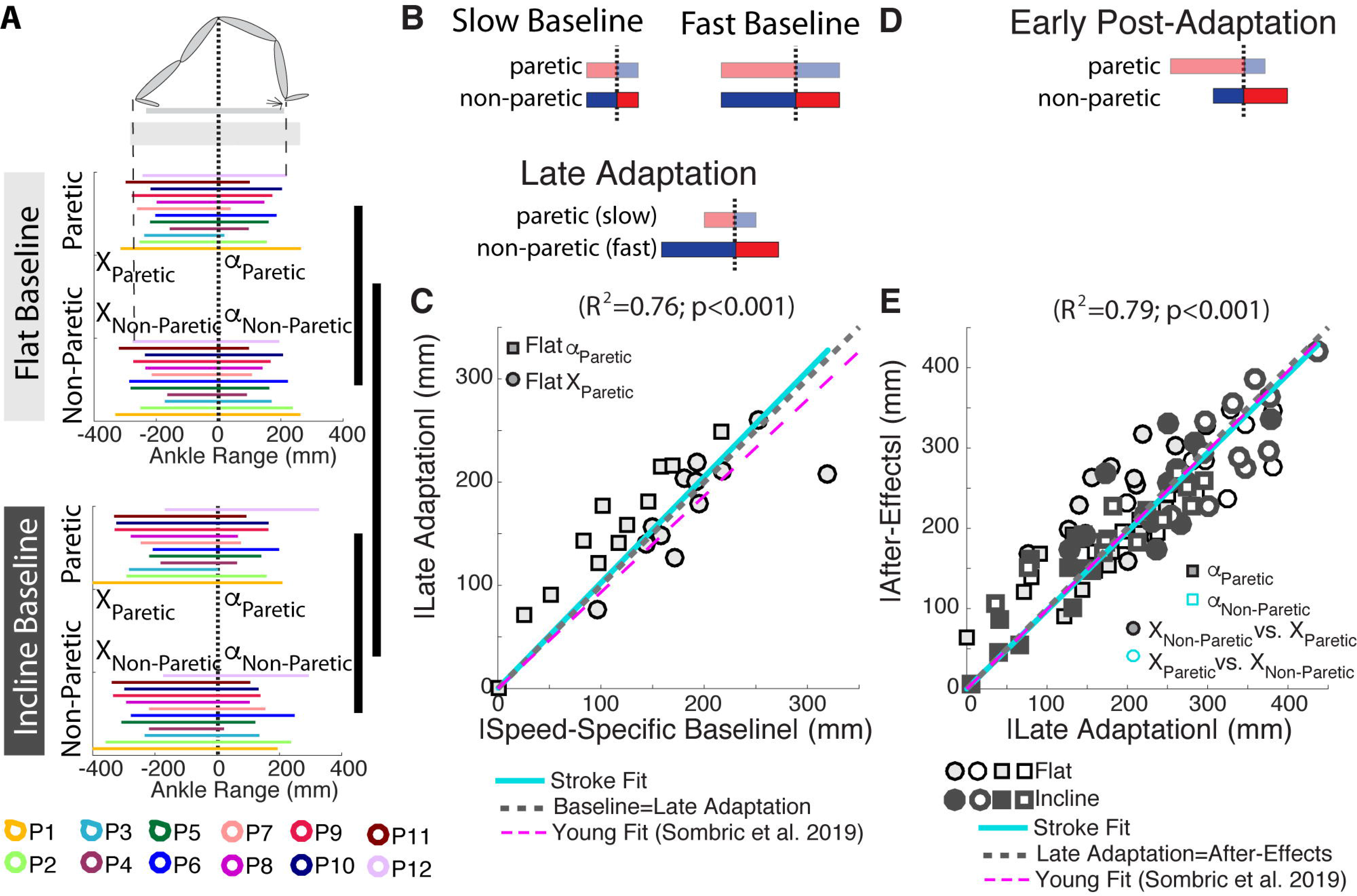
Leg orientation Adaptation and After-Effects. **(A)** Leg orientations are depicted for individual subjects (as indicated with different colors) in both the flat and incline conditions. Note that subjects orient their legs about their bodies differently and that leg orientations are based on slope. Thick vertical black lines indicated a significant effect of leg (i.e., paretic or non-paretic) and slope (i.e., flat or incline) on trailing leg positions. **(B)** Schematic of the slow and fast (predicted) baseline behavior for the paretic and non-paretic leg orientations, respectively. The speed-specific leg orientations were regained during Late Adaptation. **(C)** The similarity between leg orientations across the speed-specific Baseline and Late Adaptation epochs is illustrated by the significant regression (solid cyan line; |*y|* = *a*∗|*x|*, 95% confidence interval for *a* = [0.92, 1.13]). Recall that a slow Baseline was only collected in the flat session, thus only the slow Baseline and Late Adaptation for the paretic leg (which walked slow during Adaptation) are shown. Note that the regression line closely overlaps with the idealized situation in which baseline and late adaptation values are identical (dashed gray line; i.e., *y* = *x*) and the behavior of young, healthy adults (Sombric et al. 2019, dashed magenta line). **(D)** Schematic of the leg orientations during early Post-Adaptation. The forward leg positions are ipsilaterally and the trailing leg positions are contralaterally maintained from split-to-tied walking. **(E)** The ipsilateral and contralateral similarity between α and *X*, respectively, across the Late Adaptation and early Post-Adaptation epochs is quantified with a significant correlation (solid cyan line; |*y|* = *a*∗|*x|*, 95% confidence interval for *a* = [0.94, 1.02]). The idealized situation in which Late Adaptation and early Post-Adaptation values are identical (dashed gray line; i.e., *y* = *x*) and the behavior of young, healthy adults ((Sombric et al. 2019, dashed magenta line) are presented as a reference.

Moreover, we found that the leg orientations achieved during Late Adaptation were predictive of subjects’ Post-Adaptation behavior (Figure 4D). Specifically, the leading leg’s orientations were similar before and after removal of the split-belt perturbation (i.e., Late Adaptation α_Paretic_=Post-Adaptation α_Paretic_ and vice versa) whereas the trailing legs’ orientations were swapped between the legs (i.e. Late Adaptation X_Paretic_ = Post-Adaptation X_Non-Paretic_ and vice versa). This is supported by the significant relationship between Late Adaptation and Post-Adaptation leg orientations observed when individual subjects’ values for each leg and both sloped sessions are regressed (Figure 4E; solid cyan line; |y|=a*|x|; 95% confidence interval of a=[0.94, 1.02], R^2^=0.79, p<0.001). We also show as a reference, the relation between (recorded) Late Adaptation and (predicted) Post-Adaptation leg orientation values for both legs and both sloped conditions in young, intact individuals (magenta dashed line; (|y|=a*|x|; 95% confidence interval of a=[0.95, 1.03], R^2^=0.78, p<0.001). Note the similarity between the intact and lesioned behavior (cyan vs. magenta lines). Similar to the intact motor system, the lesioned motor system is able to recover speed and slope-specific leg orientations during Late Adaptation, which predict after-effects during Post-Adaptation.

### Larger after-effects of propulsion forces split-belt incline walking

Sloped walking influenced the extent of recalibration of the non-paretic propulsion forces. Figure 5A shows that propulsion forces were altered during the Adaptation epochs. These data are plotted relative to Baseline propulsion forces, which were larger in the incline condition and the non-paretic leg for both sloped conditions (Figure 5B: p_Individual_=0.007, p_Slope_<0.0001, p_Leg_=0.040, p_Slope#Leg_=0.43). Note that subjects were closer to generating Baseline-like propulsion forces during Late Adaptation in the incline session compared to the flat session for both legs, resulting in larger Late Adaptation paretic propulsion forces in the incline session (Figure 5C). Even though the Late Adaptation behavior was different across sessions (Figure 5C; non-paretic propulsion: p=0.032, paretic propulsion: p=0.015), the changes in propulsion forces from early to late Adaptation were similar across sloped conditions (Figure 5D; non-paretic propulsion: p=0.92, paretic propulsion: p=0.33). While paretic propulsion After-Effects are similar in either sloped conditions (Figure 5E, p=0.43), the non-paretic After-Effects are larger in magnitude following incline adaptation (p=0.015). Note that the paretic propulsion forces change the most during Adaptation (Figure 5C), whereas the non-paretic propulsion forces are the ones exhibiting after-effects during Post-Adaptation (Figure 5E). In summary, incline walking results in augmented paretic propulsion forces during Adaptation and reduced non-paretic propulsion force After-Effects.

**Figure 5:**
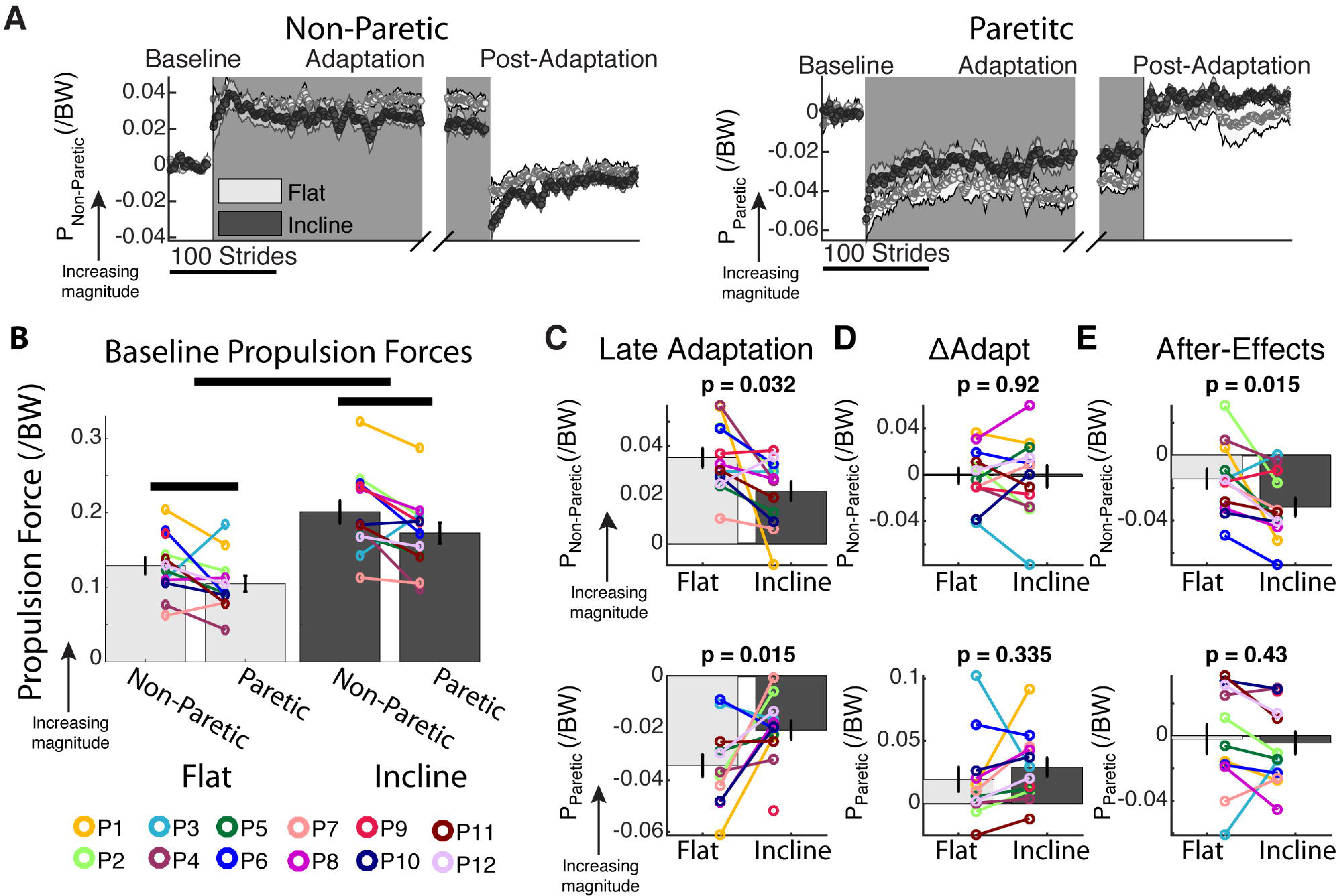
Propulsion force Adaptation and After-Effects. **(A)** Stride-by-stride time courses of propulsion forces of the non-paretic (top panel) and paretic leg (bottom panel) are shown during self-selected Baseline, Adaptation, and Post-Adaptation. Each data point represents the average of 5 consecutive strides and shaded regions indicate the standard error for each group. For display purposes only, we include stride values during Post-Adaptation that were computed with a minimum of 10 subjects. **(B-E)** We display group average values for propulsion force outcome measures ± standard errors. Individual subjects are represented with colored dots connected with lines. **(B)** Baseline: Thick horizontal black lines indicated that there is a significant effect of leg (i.e., paretic or non-paretic) and slope (i.e., flat or incline) on propulsion forces. On average, stroke subjects generate larger propulsion forces with their non-paretic leg, and they generate larger propulsion forces with both legs when walking incline. However, some individual stroke subjects generate larger propulsion forces with their paretic than their non-paretic leg. **(C)** ΔAdapt: Propulsion forces were similarly modulated during the Adaptation epoch for both sloped conditions. **(D)** Late Adaptation: Stroke subjects were closer to their baseline propulsion forces in the incline than the flat sessions. Moreover, baseline propulsion forces in the incline session were larger than the flat session (Figure 2C). Taken together, these results suggest that stroke subjects are forced to propel more during incline split-belt walking with both legs compared to flat split-belt walking. **(E)** After-Effects: Even though both sloped sessions did not change the extent of propulsion force adaptation (ΔAdapt), slope influenced the After-Effects for the non-paretic leg, but not the paretic leg.

## Discussion

We investigated the influence of locomotor propulsion demands on motor adaptation and recalibration of gait in the asymmetric motor system. We find that survivors of a stroke adapt more during incline than flat split-belt walking. We also find that leg orientations during Adaptation are predictive of those Post-Adaptation, leading to greater step length asymmetry after-effects in the incline than flat sessions. Lastly, larger step length asymmetry after-effects resulted from shorter paretic step lengths and lower non-paretic propulsion forces during Post-Adaptation in the incline session. In summary, the ability to control leg orientation to meet speed and force demands during split-belt walking is maintained post-stroke, which can be exploited for designing effective gait rehabilitation interventions.

### Post-stroke gait adapts more in response to larger propulsion demands

We find that survivors of a stroke behave similarly to young, intact adults in their response to sloped split-belt walking. Specifically, survivors of a stroke are able to augment their propulsion forces and adjust their leg orientations in response to incline split-belt walking as observed in young, healthy adults (Sombric et al. 2019). This is consistent with previous literature indicating that patients in the chronic phase post-stroke can modulate their gait in response to task demands (Kesar et al. 2011, 2014; Reisman et al. 2013; Awad et al. 2014; Hsiao et al. 2015, 2016b). Additionally, we observe that survivors of a stroke recover speed and slope-specific paretic leg orientations in the flat session. We speculate that the same would have been observed in both legs and sloped conditions like young adults (Sombric et al. 2019). This is a reasonable expectation given that survivors of a stroke exhibit similar control of leg orientations to young adults during Late Adaptation and early Post-Adaptation for both legs and sloped conditions. Thus, our results provide further evidence that steady state split-belt walking can be predicted from Baseline walking.

It has been previously suggested that patients’ baseline gait asymmetries determine motor behavior at steady state split-belt walking (Reisman et al. 2007; Malone and Bastian 2014). Our results provide new insight that survivors of a stroke recover their baseline asymmetry in the incline, but not in the flat condition. Thus, it is not baseline gait asymmetry, but kinetic demands that might govern patients’ motor patterns. More specifically, our results suggest that survivors of a stroke aim to recover the baseline leg orientations for the kinetic demands for each leg in the split condition, as observed in young adults (Sombric et al. 2019). While our observed forward leg orientations are suited for harnessing energy from the treadmill, the trailing leg orientations are not (Sánchez et al. 2019)(additional file 1). Thus, there might be some other factors such as stability (Buurke et al. 2018) or metabolic energy (Gordon et al. 2009) regulating leg orientation in walking. In summary, the forces generated to propel one’s body forward constitute an important control variable regulating the adaptation of movements in the intact and asymmetric motor systems.

### Bilateral adaptation in survivors of a stroke contrasts unilateral adaptation in young adults

Survivors of a stroke recruit both legs in order to adapt their gait, whereas young adults primarily adapt one leg. Notably, while survivors of a stroke adapt both the paretic (slow belt) and nonparetic (fast belt) step lengths, whereas we previously found that young individual predominantly adjust the fast belt step length (Sombric et al. 2019). This could be because survivors of a stroke may require more repetitions in the altered environment to recover their baseline leg orientation with their paretic leg, whereas intact subjects can do so immediately after the split condition is introduced. Alternatively, it could be that the larger neural coupling post-stroke (Kloter et al. 2011) enhances bilateral adaptation. Regarding post-adaptation, paretic after-effects are only observed in the incline condition. More specifically, paretic step lengths become longer than in baseline walking, which is beneficial for survivors of a stroke who normally take short paretic step lengths (Balasubramanian et al. 2007). On the other hand, non-paretic after-effects are observed regardless of the sloped condition. This is atypical since the non-paretic leg walked fast in the split condition and young adults only exhibit after-effects in the leg that walked slow (Sombric et al. 2019). Specifically, survivors of a stroke take non-paretic step lengths shorter than those taken during Baseline perhaps as a strategy to recover balance (e.g., Eng et al., 1994), which is challenged upon removal of the split condition (Buurke et al. 2018; Iturralde and Torres-Oviedo 2019). In summary, survivors of a stroke adapt both legs during split-belt walking, but paretic step length after-effects are only observed following incline split-belt walking.

### Neurorehabilitation through reinforcement of a corrective pattern during adaptation, rather than short-lived after-effects post-adaptation

The long-term therapeutic effect of locomotor adaptation with split-belt treadmills may be due to walking with the motor demands of the split-belt task, rather than the adaptation effects observed post-adaptation. Split-belt walking has been shown to reduce long term gait asymmetry (Reisman et al. 2013; Lewek et al. 2018). However, it is unclear what aspect of split-belt walking underlies these changes. After-Effects could lead to motor improvements (Bastian 2008) such as reduced gait asymmetry immediately (Reisman et al. 2007; Choi et al. 2009). However, these after-effects are short lived and decrease as individuals experience multiple days of practicing the split-belt condition (Larish et al. 1988; Sombric et al. 2017; Leech et al. 2018). It is known that regular treadmill walking cannot modify gait asymmetries post-stroke (Silver et al. 2000; Kautz et al. 2005; Den Otter et al. 2006), suggesting that the specific motor demands of the split-belt task might be important for neurorehabilitation. For example, we observe that the split condition forces survivors of a stroke to take longer paretic step lengths and generate greater paretic propulsion forces. Perhaps practice of these gait features during multiple exposure to the split situation might lead to long term changes in gait symmetry. It is also possible that the strenuous nature of split-belt walking increases neural plasticity, as shown with other high-intensity exercises (Andrews et al. 2019). Thus, incline split-belt walking may be beneficial not only for inducing greater paretic propulsion, but also because it is more demanding than level walking (Johnson et al. 2002). Lastly, it is also possible that the initial disruption of step length asymmetry is fundamental (Marchal-Crespo et al. 2014) for patients to start exploring new locomotor patterns that could converge to more metabolically efficient gait than their baseline walking pattern (Selinger et al. 2015; Sánchez et al. 2019). In summary, the long term benefit of split-belt walking may originate from practicing motor patterns specific to split-belt walking, rather than reinforcement of those observed during post-adaptation.

### Clinical implications

Split-belt walking has been shown to induce long term changes that could improve the mobility of those who have had a stroke (Reisman et al. 2013; Betschart et al. 2018; Lewek et al. 2018). However, it is unclear which leg should be placed on the slow and fast belt during the split condition (Reisman et al. 2007, 2009; Malone and Bastian 2014; Finley et al. 2015). For instance, one may consider that the paretic leg should walk on the slow belt to force survivors of a stroke to use it more, as a form of “constrained use therapy” (e.g., Kwakkel et al. 2015). On the other hand, our results here and in a previous study (Sombric et al. 2019) indicate that placing the paretic leg on the fast belt would force subjects to augment their paretic propulsion forces and lengthen their paretic steps during split-belt walking, which could be beneficial. Future studies are needed to determine if survivors of a stroke could actually augment their paretic propulsion forces during split-belt walking, as observed in the fast leg of young adults (Sombric et al. 2019). This remains an open question given that we observed limited changes in paretic propulsion post-adaptation compared to those reported in controls (Sombric et al. 2019). In sum, our study provides greater understanding of the motor demands associated to the split-belt task, which could be harnessed for gait neurorehabilitation.

Incline split-belt training may be a promising way to augment locomotor and adaptation and recalibration in the lesioned motor system. Not everyone who has had a stroke re-learns to walk symmetrically following several weeks of flat split-belt training (Reisman et al. 2013; Betschart et al. 2018; Lewek et al. 2018). Thus, it is clinically relevant to explore alternative strategies to augment adaptation in survivors of stroke other than increasing the speed difference (Yokoyama et al. 2018) since not all patients can walk with large speed differences. While this work indicates that adaptation can be augmented in patients, previous work indicates that overground walking post-stroke is most improved following decline, rather than incline, interventions (Carda et al. 2013). Thus, future studies are needed to determine if the augmented adaptation in the incline environment transfers to flat overground walking.

## Conclusion

We investigated the influence of propulsion demands during walking on the locomotor adaptation and recalibration in the asymmetric motor system. We found that survivors of a stroke adapt more when propulsion demands are increased. Like intact subjects, after-effects can be predicted by each patient’s leg orientations achieved during split-belt walking, which in turn were predicted by subject-specific leg orientations during baseline walking. These results have two implications. First, these findings indicate that survivors of a stroke are able to adjust their movements to meet kinetic demands imposed by the walking condition. Second, our results suggest baseline motor features can be predictive of the individual’s ability to adjust their movements. In summary our study provides valuable insights into the mechanisms underlying locomotor adaptation induced by split-belt walking, which could be exploited for designing effective gait rehabilitation interventions post-stroke.

## List of Abbreviations

SL: step length
α: leading leg position relative to the hip at leading leg heel strike
X: trailing leg position relative to the hip at leading leg heel strike
GRF: ground reaction forces
AP: anterior-posterior
P_Pareitc_: paretic peak propulsion forces
P_Non-Pareitc_: non-paretic peak propulsion forces

## Declarations

### Ethics approval and consent to participate

Written and informed consent was obtained from all participants prior to participation. The University of Pittsburgh Institutional Review Board approved the used experimental protocol.

### Consent for Publication

No applicable.

### Availability of data and materials

The datasets used and/or analyzed during the current study are available from the corresponding author on reasonable request.

### Authors’ contributions

G.T.-O. and C.S. were involved with the conception and design of the work. C.S. collected and analyzed the data. C.S. and G.T.-O. interpreted the results. C.S. drafted the manuscript, which was carefully revised by all authors. The final version of the manuscript has been approved by all the authors.

## Acknowledgements

The authors acknowledge the valuable input from Pablo Iturralde and Digna de Kam.

