## additional file 1 for "Augmenting propulsion demands during split-belt walking increases locomotor adaptation in the asymmetric motor system"

### **Additional Files**

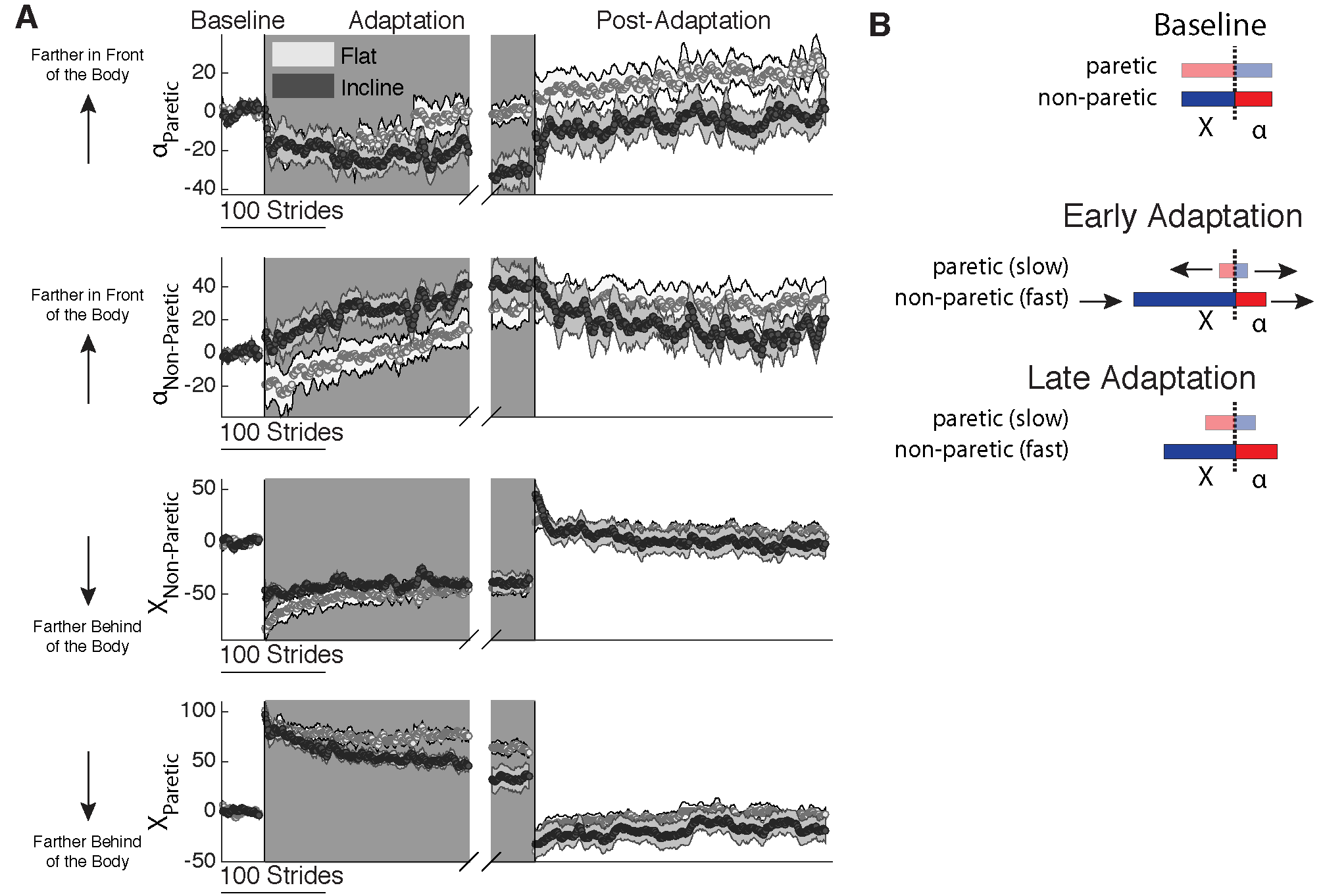

**Additional File 1: Leg Position Adaptation and After-Effects in the Asymmetric Motor System| (A)** Stride-by-stride time courses of leg positions (α and X) for the non-paretic and paretic leg are shown during self-selected Baseline, Adaptation, and Post-Adaptation. Each data point represents the average of 5 consecutive strides and shaded regions indicate the standard error for each group. The beginning and Late Adaptation group average behavior are shown for the Adaptation epoch. For display purposes only, we include stride values during Post-Adaptation that were computed with a minimum of 10 subjects. **(B)** Schematic of the self-selected Baseline, early Adaptation, and late Adaptation behavior for the paretic and non-paretic leg orientations, respectively. Note that there is a general forward movement of the leg position of the non-paretic leg, but the paretic leg increases both the leading and trailing positions.
